# The role of posterior ventral stream areas for viewpoint-invariant object recognition

**DOI:** 10.1101/2021.07.07.451407

**Authors:** Sophia Nestmann, Hans-Otto Karnath, Johannes Rennig

**Author notes:** Address for correspondence:* Hans-Otto Karnath, Center of Neurology, University of Tübingen, D-72076 Tübingen, Germany.

## Abstract

Object constancy is one of the most crucial mechanisms of the human visual system enabling viewpoint invariant object recognition. However, the neuronal foundations of object constancy are widely unknown. Research has shown that the ventral visual stream is involved in processing of various kinds of object stimuli and that several regions along the ventral stream are possibly sensitive to the orientation of an object in space. To systematically address the question of viewpoint sensitive object perception, we conducted a study with stroke patients as well as an fMRI experiment with healthy participants applying object stimuli in several spatial orientations, for example in typical and atypical viewing conditions. In the fMRI experiment, we found stronger BOLD signals and above-chance classification accuracies for objects presented in atypical viewing conditions in fusiform face sensitive and lateral occipito-temporal object preferring areas. In the behavioral patient study, we observed that lesions of the right fusiform gyrus were associated with lower performance in object recognition for atypical views. The complementary results from both experiments emphasize the contributions of fusiform and lateral-occipital areas to visual object constancy and indicate that visual object constancy is particularly enabled through increased neuronal activity and specific activation patterns for objects in demanding viewing conditions.

## Introduction

Visual constancy is a crucial mechanism of the human visual system that allows the perception of familiar objects regardless of changes in perspective, distance, lighting, or size (Brunswik, 1934; Hebb, 1958; Emmert, 1881; Fitzpatrick et al., 1982). How the human brain solves such complex object recognition is still under debate. Behavioral data have suggested a viewpoint-dependent representation of familiar objects (e.g., Tarr & Pinker, 1989; Ullman, 1989; Bülthoff & Edelman, 1992; Tarr, 1995; Tarr et al., 1998; Andresen et al., 2009), but also viewpoint-invariant representations based on an object’s subcomponents (e.g., Biederman, 1987; Biederman & Gerhardstein, 1993). Jolicoeur (1990) proposed a dual-system theory uniting both approaches suggesting two functionally separate systems working in parallel and facilitating object recognition: a mental-rotation system, which rotates the perceived stimulus to the vertical for comparisons with memory representations, and a feature-based system, which is orientation invariant. Today, much research focuses on computer vision, aiming at modelling cognitive processes and cortical mechanisms in order to solve questions on object recognition (e.g., Kreiman & Serre, 2020).

Numerous lesion and functional neuroimaging studies have demonstrated the crucial role of the ventral visual stream in object recognition (Ungerleider & Mishkin, 1982; Goodale et al., 1991; Milner et al., 1991; Goodale & Milner, 1992; James et al., 2003; Milner & Goodale, 2008; Karnath et al., 2009). However, how specific brain areas along the ventral stream solve invariant object recognition is still widely unknown: Logothetis & Pauls (1995) showed that inferotemporal cortices of nonhuman primates responded selectively to different views of familiar objects. In an fMRI experiment, Vuilleumier et al. (2002) observed that the right fusiform gyrus was sensitive to changes in viewpoint during the presentation of three-dimensional objects. Pourtois et al. (2005) demonstrated that the lateral fusiform cortex responds selectively to viewpoints of familiar faces. However, other fMRI studies revealed that objects presented at different viewpoints have similar univariate signal characteristics suggesting viewpoint-invariant representations in the ventral stream (James et al., 2002; Valyear et al., 2006). Eger et al. (2008) showed that a classifier was able to categorize object stimuli from brain activity in the lateral occipital complex (LOC, Grill-Spector et al., 2001) and early visual cortex across different viewpoints. This finding indicated that objects might be represented in a viewpoint invariant manner in these brain areas.

To this point, previous research provided ambiguous evidence of viewpoint-invariant object processing in the ventral visual stream and the specific role of object-sensitive areas, like the LOC, for visual object constancy. In a first study, we defined three regions of interest (ROIs) along the ventral stream and compared BOLD signals in these areas during the processing of objects shown from different views. The objects were presented in usual (*canonical*) and unusual (*non-canonical*) views as well as in *frontal* and *in-depth* rotated presentations. We aimed to detect differences in signal strength in one or more of our ventral ROIs between objects presented in *canonical* and *non-canonical* views (Nestmann et al., 2021) as well as between *frontal* and *in-depth* views. In a machine learning approach, we trained a classifier with *canonical/non-canonical* and *frontal/in-depth* rotated object views and explored the viewpoint specificity in the three ventral ROIs. In a second, behavioral study with stroke patients, we assessed the effects of lesions to the ventral stream to viewpoint-invariant objects perception.

## Methods

### A. fMRI study

Twenty healthy volunteers (mean age = 25.7 years; *SD* = 3.9 years, 10 female, 1 left-handed) with no history of neurological or psychiatric disorders participated in the study after giving written informed consent. The experiment was approved by the ethics committee of the medical faculty of the University of Tübingen and conducted in accordance with the declaration of Helsinki. All participants had normal or corrected to normal vision.

Imaging was performed using a 3T Siemens Magnetom TrioTim MRI system (Siemens AG, Erlangen, Germany) equipped with a 64-channel head coil. During a scanning session, participants performed two independent fMRI experiments. The first fMRI experiment was an event-related design. Stimuli were presented for 300 ms with an inter-stimulus interval of 1700 ms. The events were ordered in an optimal rapid event-related design specified by optseq2 (Dale, 1999; https://surfer.nmr.mgh.harvard.edu/optseq). For the second fMRI experiment (localizer task), we applied a block design. Stimuli were presented using Matlab (The Mathworks, Inc., Natick, MA, USA) and the Psychophysics Toolbox (Brainard, 1997; Pelli, 1997) and projected onto an MR compatible screen placed behind the scanner bore which could be viewed by the participants via a mirror mounted on the head coil. Behavioral responses were collected using a fiber-optic button response pad (Current Designs, Haverford, PA, USA).

In the first fMRI experiment, participants were presented with computer-generated object stimuli in different viewing conditions that were taken from the Object Databank (http://wiki.cnbc.cmu.edu/Objects) and from 3dcadbrowser (https://www.3dcadbrowser.com). As previous studies used different definitions of ‘canonical’ and ‘non-canonical’ viewing conditions varying from ‘frontal’ views (e.g., Best, 1917; Solms et al., 1988; Friedman-Hill et al., 1995; Turnbull et al., 1995, 1997a; Karnath et al., 2000) to ‘in-depth rotations’ (e.g., Terhune et al., 2005; Landau et al., 2006; Schendan & Stern, 2008), we created a systematic stimulus set that consisted of ‘frontal’ and ‘in-depth rotated’ views both as ‘canonical’ and ‘non-canonical’ conditions (Fig. 1a). This allowed us to address potential differences between neuronal processing of the views. The experiment was carried out in four blocks, each consisting of 144 stimuli (36 of each viewing condition) presented in pseudo-random order (duration of each block: about 5.5 minutes). The stimuli were presented in grayscale and were normalized for luminance and spatial frequencies and controlled for semantic frequency between the four experimental conditions. The objects were presented at a size of 5° visual angles. To ensure that participants were paying attention to the object stimuli we conducted a task independent from the objective of our study, in which we asked participants to classify the presented stimuli as either ‘made from metal’ or ‘not made from metal’ by button press. In a recent study (Nestmann et al., 2021), it was demonstrated that this particular task does not bias the results of our experiments.

**Figure 1.**
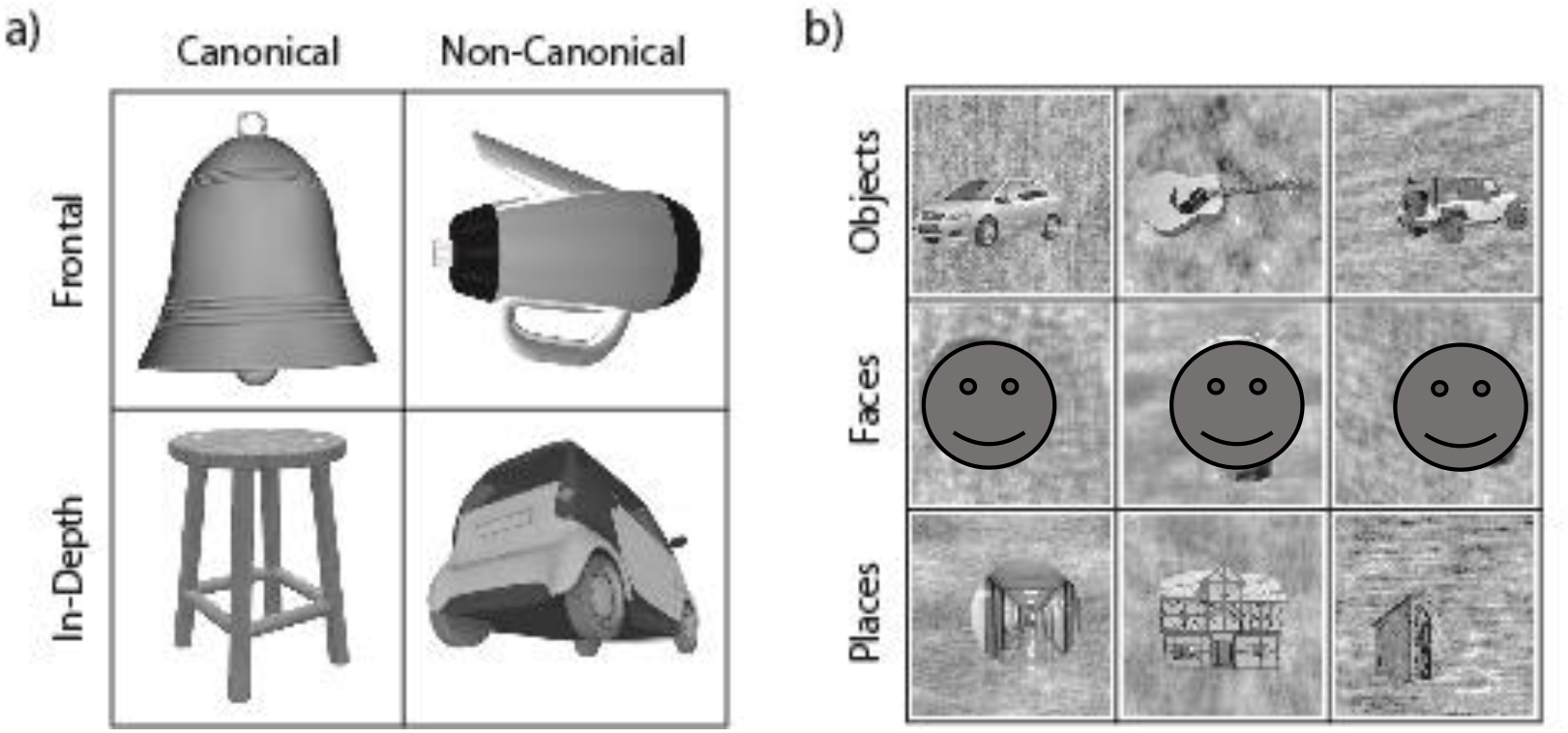
Sets of stimuli used in the two fMRI experiments. **a)** In the first experiment, objects from four different categories were presented. These included objects in ‘frontal’ and ‘in-depth rotated’ views both as ‘canonical’ and ‘non-canonical’ conditions. **b)** In the localizer task, we used stimuli of objects, faces and places from the study by Stigliani et al. (2015). [Due to the bioRxiv policy stimuli of faces have been covered.]

The second fMRI experiment was a localizer task. Here, we determined individual ROIs sensitive for objects, faces and places. In a block design, participants were presented with images of objects, faces and places taken from Stigliani et al. (2015) at 5.2° visual angles in a one-back task (cf. Fig. 1b). They were instructed to indicate by button press, when one image was directly repeated. Stimuli were presented in two blocks (duration of each block about 6 minutes) with 30 sections. Each section consisted of eight stimuli from only one category. Images were presented for 0.5 seconds with an inter-stimulus interval of 1 second. Consecutive sections were separated by an interval of 2 seconds.

#### MRI data acquisition

Functional images were acquired using multiband echo-planar-imaging (EPI) sequences. The first three participants were scanned using the following scanning parameters: TR = 2000 ms, TE = 33 ms, flip angle = 58°, FOV = 1728 × 1728 mm^2^, 69 slices, voxel size = 2 × 2 × 2 mm^3^, multiband factor = 3. All remaining participants were scanned with parameters for multiband EPI sequences from the HCP (Moeller et al., 2010): TR = 1000 ms, TE = 37 ms, flip angle = 52°, FOV = 1872 × 1872 mm^2^, 72 slices, voxel size = 2 × 2 × 2 mm^3^. Single band reference images (TR = 1000 ms, TE = 37 ms, flip angle = 52°, FOV = 1872 × 1872 mm^2^, 72 slices, voxel size = 2 × 2 × 2 mm^3^) were collected before each functional run. T1-weighted anatomical scans (TR = 2280 s, 176 slices, voxel size = 1.0 × 1.0 × 1.0 mm^3^; FOV = 256 × 256 mm^2^, TE = 3.03 ms; flip angle = 8°) were collected at the end of the experimental session.

#### fMRI data analysis

Data pre-processing and model estimation were performed using SPM12 (http://www.fil.ion.ucl.ac.uk/spm). Functional images were realigned to each participants’ first image, aligned to the AC-PC axis and slice-time corrected. The original single-band image was then co-registered to the pre-processed functional images and the anatomical image was co-registered to the single-band image. The resolution of the single-band image was up-sampled before the anatomical image was aligned to it. Functional images were smoothed with a 4 mm FWHM Gaussian kernel. Time series of hemodynamic activation were modelled based on the canonical hemodynamic response function (HRF) as implemented in SPM12. Low-frequency noise was eliminated with a high-pass filter of 128 Hz. Correction for temporal autocorrelation was performed using an autoregressive AR(1) process. Movement parameters (roll, pitch, yaw; linear movement into x-, y-, z-directions) estimated in the realignment were included as regressors of no interest. To avoid bias associated with spatial normalization, analyses were conducted in native space. For the first fMRI experiment, the experimental regressors consisted of the four experimental conditions: *canonical rotated in-depth view, canonical frontal view, non-canonical in-depth view* and *non-canonical frontal view*. For the localizer task we included the regressors *objects, faces* and *places*.

#### Region of interest (ROI) analysis

We included the LOC as a generally object-sensitive area (e.g., Grill-Spector et al., 2000; Kourtzi & Kanwisher, 2000; Grill-Spector, 2003), the fusiform face area (FFA; Puce et al., 1995; Kanwisher et al., 1997) since it was demonstrated that the fusiform gyrus is significantly involved in processing of objects from different viewpoints (Kosslyn et al., 1994; Sugio et al., 1999; Vuilleumier et al., 2002), and the parahippocampal place area (PPA) which has previously been associated with the processing of complex visual stimuli, like scenes (e.g., Epstein & Kanwisher, 1998). Using this area allowed us to explore the role of more anterior located brain areas during the processing of objects in unusual views.

Anatomical ROIs were created applying Freesurfer’s cortical reconstruction routine (Dale et al., 1999; Fischl et al., 2002) and the Destrieux atlas (Destrieux et al., 2010) for each subject. We used Freesurfer Labels 11158/12158, 11160/12160, 11159/12159 (anterior, middle and superior occipital cortex, transverse and lunate sulcus) to create an anatomical ROI of the lateral occipito-temporal cortex. We selected Freesurfer Label 11121/12121 for the fusiform gyrus and Label 11123/12123 for the parahippocampal gyrus. In a next step, we identified individual object-sensitive voxels comparing blocks of object presentations to blocks of faces and places. To identify face-sensitive voxels we compared blocks of face presentations to blocks of objects and places. We identified place-sensitive contrasting blocks of place images to blocks of objects and faces. The voxel-level threshold was set to *p* < 0.05 (uncorr.) without cluster threshold. Objects ROIs (corresponding to LOC) were created as the intersection between the lateral occipito-temporal cortex and object-sensitive voxels, faces ROIs (corresponding to FFA) came from the intersection of the fusiform gyrus and face-sensitive voxels and places ROIs (corresponding to PPA) were created as the intersection between parahippocampal cortex and place sensitive voxels. ROIs for objects, faces and places were identified in all 20 participants in both hemispheres. Examples for structural and functional ROIs are presented in Figure 2. ROI characteristics are summarized in Table 1.

**Table 1.**
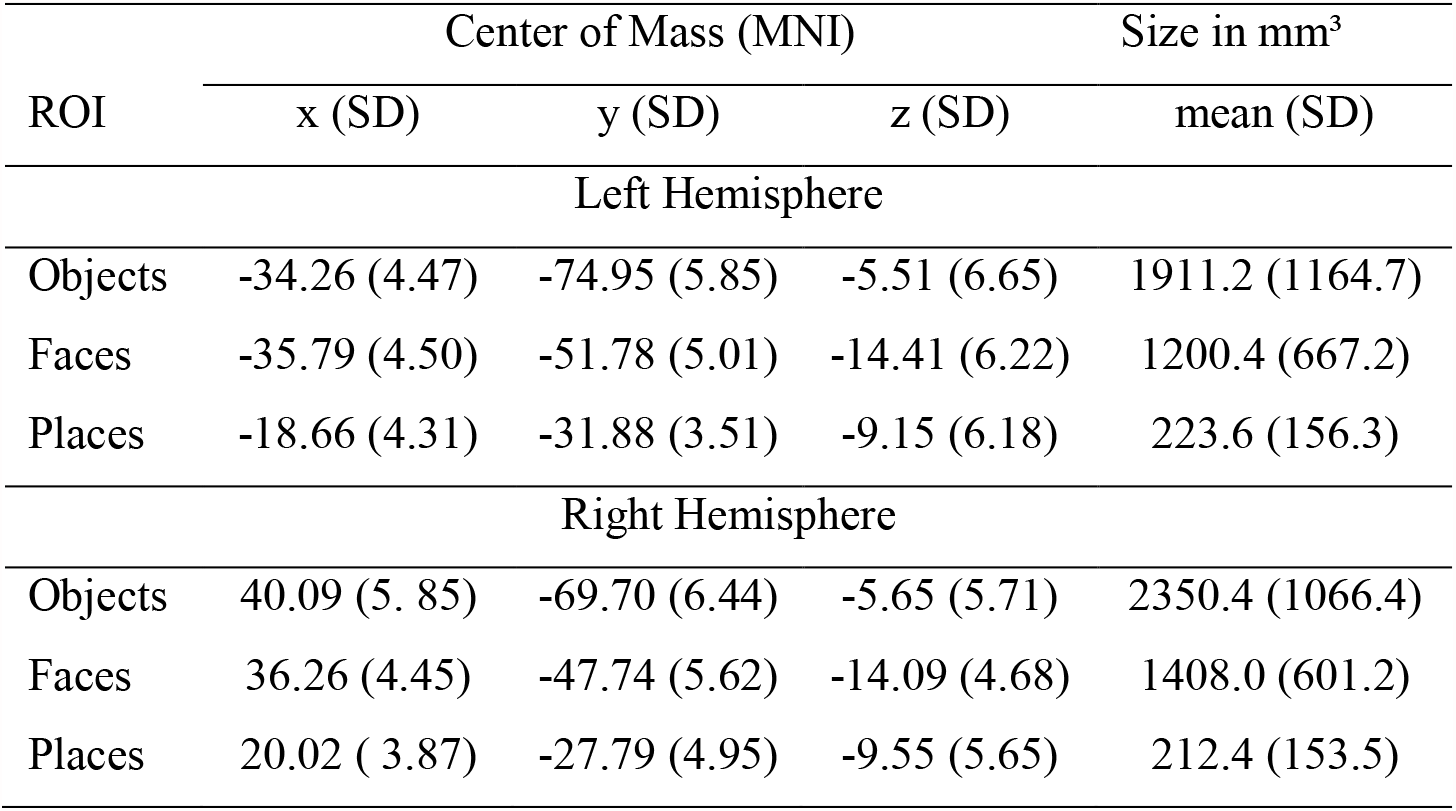
ROI locations and size.

| ROI | Center of Mass (MNI) |  |  | Size in mm<sup>3</sup> |
| --- | --- | --- | --- | --- |
|  | x (SD) | y (SD) | z (SD) | mean (SD) |
| Left Hemisphere |  |  |  |  |
| Objects | -34.26 (4.47) | -74.95 (5.85) | -5.51 (6.65) | 1911.2 (1164.7) |
| Faces | -35.79 (4.50) | -51.78 (5.01) | -14.41 (6.22) | 1200.4 (667.2) |
| Places | -18.66 (4.31) | -31.88 (3.51) | -9.15 (6.18) | 223.6 (156.3) |
| Right Hemisphere |  |  |  |  |
| Objects | 40.09 (5.85) | -69.70 (6.44) | -5.65 (5.71) | 2350.4 (1066.4) |
| Faces | 36.26 (4.45) | -47.74 (5.62) | -14.09 (4.68) | 1408.0 (601.2) |
| Places | 20.02 (3.87) | -27.79 (4.95) | -9.55 (5.65) | 212.4 (153.5) |

**Fig. 2.**
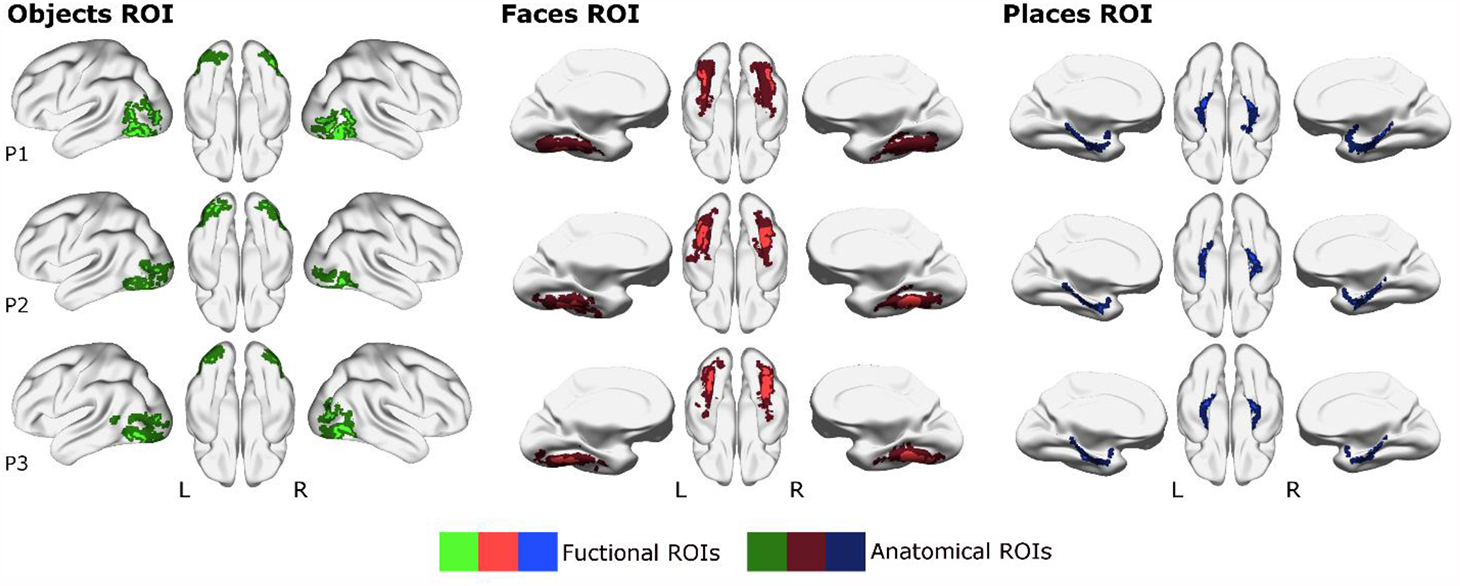
Examples of the ROIs used. Example ROIs from three subjects are presented in standard MNI space on a surface version of the ch2 brain using the BrainNet viewer (Xia et al., 2013). Individual anatomical lateral occipito-temporal, fusiform and parahippocampal ROIs were created applying Freesurfer’s cortical reconstruction routine (Dale et al., 1999; Fischl et al., 2002) and the Destrieux atlas (Destrieux et al., 2010). From these ROIs, we then identified voxels that selectively responded to objects, faces and places. objects ROI, faces ROI and places ROI were created as the respective intersections. Percent signal change values and beta-coefficients were extracted from these individual ROIs and used for univariate and multivariate analyses.

We used MarsBar (http://marsbar.sourceforge.net) to extract the mean percent signal change from individual ventral objects, faces and places ROIs for all four experimental conditions. For statistical data analysis, we applied linear mixed effect models using R’s *lme4* and *lmerTest* packages. Model estimation was done using Restricted Maximum Likelihood (REML) estimation. Statistical significance was assessed using the *Anova* function provided by the *car* package. Estimated marginal means were calculated by the *emmeans* function, effect sizes by the *eff_size* function provided by the *emmeans* package (Searle et al., 1980).

#### Multivoxel Pattern Analysis (MVPA)

A multivariate analysis (multivoxel pattern analysis, MVPA) can quantify neuronal representational similarities between experimental conditions (Haxby et al., 2001; Haynes & Rees, 2006) beyond comparing the univariate signal strength. We applied the method proposed by Mumford et al. (2012): for every participant we calculated beta images for each experimental trial by running a general linear model with a regressor for the respective trial as well as another regressor for all other trials. Images were not smoothed and no high-pass filtering in the statistical model applied. Beta values of voxels from each ROI were then transformed into vectors and used as features for training and testing support vector machines (SVM). The *R* package *e1071* was used to train and test SVMs.

With this analysis we aimed at exploring voxel pattern responses for canonical and non-canonical object views in object, face and place sensitive areas along the ventral visual pathway. Per participant and hemisphere, we selected beta values for every experimental trial from the fMRI experiment (canonical and non-canonical object views) from every voxel of the previously defined objects, faces and places ROIs. Each experimental trial was treated as an observation and each voxel as a feature for the machine learning model. We randomly split the data into a training set (80% of trials) and a test set (20% of trials) and trained an SVM with a radial basis kernel with the training data. We conducted a grid search using 10-fold cross validation to optimize the regularization parameter C = [0.01, 0.1, 1, 10, 100] and gamma = [0.1, 0.5, 1, 2]. Using the SVM model, we predicted from the voxel patterns of each observation of the test set (per ROI, per participant and hemisphere) if an individual trial was a canonical or a non-canonical object.

We calculated classification accuracy and predictive values per condition. The predictive value per condition was calculated as follows: number of correct classifications (condition A) / [number of correct classifications (condition A) + incorrect classification (condition A)]. The overall classification accuracy is calculated as the proportion of correctly classified trials relative to all classified trials across both conditions: [number of correct classifications (condition A) + number of correct classifications (condition B)] / [number of correct classifications (condition A)

+ number of correct classifications (condition B) + number of incorrect classifications (condition A) + number of incorrect classifications (condition B)]. Predictive values per condition correspond to sensitivity (positive predictive value) and specificity (negative predictive value) of a binary classifier. We interpret a potential difference in classification accuracies between the two conditions as noisier data in one condition compared to the other condition.

### B. Patient study

We tested a sample of 11 stroke patients with lesions in the ventral stream, a sample of 8 patients with lesions outside the ventral stream, and 12 healthy control subjects. There was no significant age difference between healthy controls, ventral patients and non-ventral patients (*F* = 0.332, *p* = 0.566). All patients had circumscribed unilateral brain lesions due to ischemic stroke or hemorrhage demonstrated by MRI (*N* = 13) or CT (*N* = 6). Lesion overlaps are shown in Figure 3. The average lesion size of ventral patients was 1502.8±1849.7 mm^3^, for non-ventral patients it was 2206.4±4265.8 mm^3^. Lesion sizes did not differ significantly between the two groups (*t* = 0.491, *p* = 0.630). The experiment was approved by the ethics committee of the medical faculty of the University of Tübingen and conducted in accordance with the declaration of Helsinki.

**Fig. 3.**
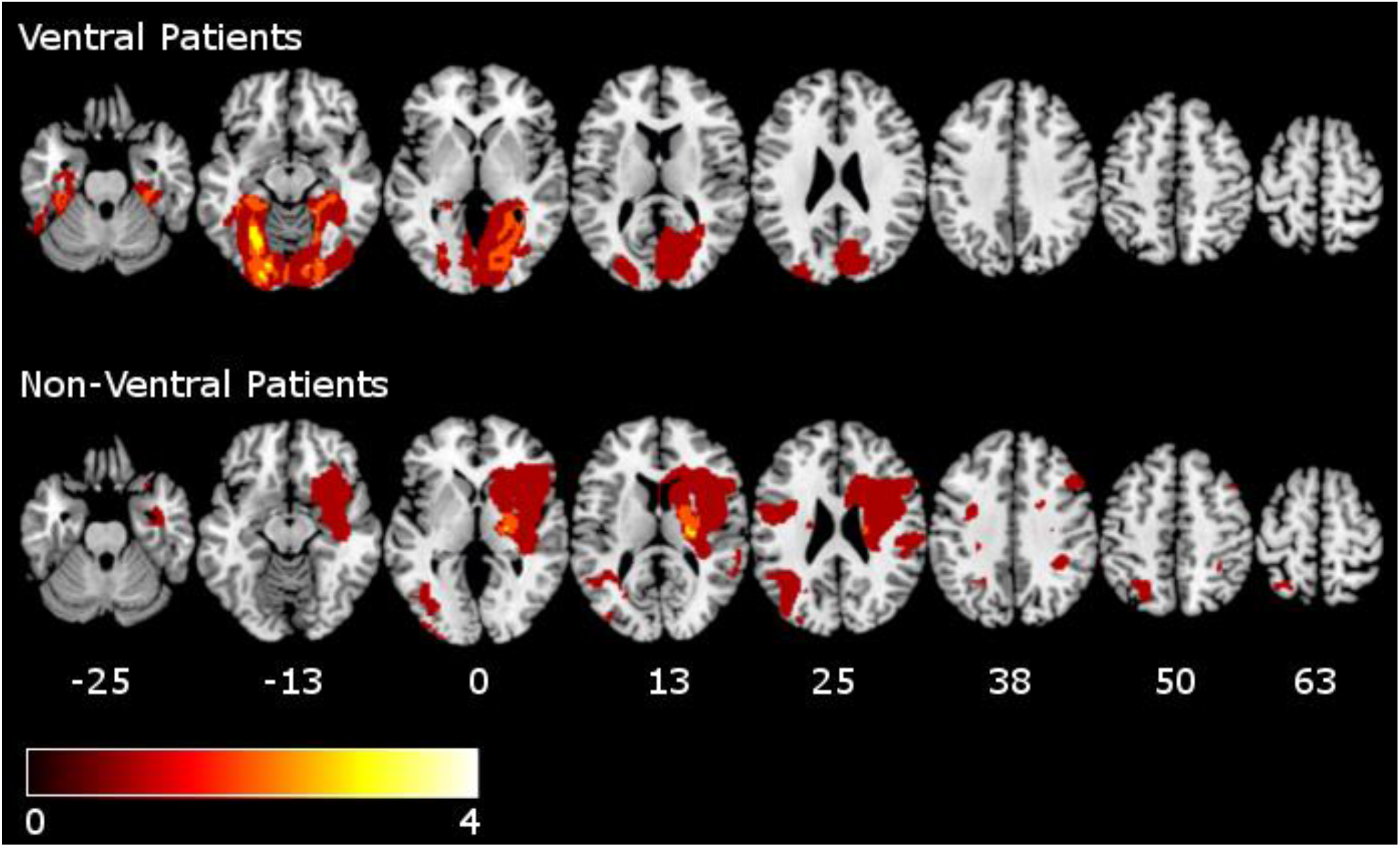
Lesion overlap of the two stroke patient groups. Lesion maps were created semi-automatically from patients’ clinical scans using the Clusterize SPM toolbox (SPM12; Clas et al., 2012; de Haan et al., 2015) and were then normalized to an age-related template with the algorithm provided by the Clinical toolbox (Rorden et al., 2012) based on SPM8 (http://fil.ion.ucl.ac.uk/spm). Normalized lesion maps of the patients are presented on the single-subject T1 MNI152 template. The vertical z coordinate of standardized MNI space is given below each slice. The color bar indicates the frequency of lesion overlap.

For acute stroke patients, the experiment was conducted at the neurological ward, all other participants were tested in an experimental room. All participants had normal or corrected to normal vision. All patients underwent neuropsychological screening for clinical object perception and naming deficits with the object perception subtests of the Visual Object and Space Perception Battery (VOSP, Warrington & James, 1991) and the short form of the Boston Naming Test (Kaplan et al., 1983) from the CERAD (Memory Clinic Basel, 2016). We also conducted tests for simultanagnosia including two figures of the overlapping figures test (Poppelreuter, 1917), ten Navon-Letters (Navon, 1977) and the Binet-Bobertag picture (Binet & Simon, 1904). We tested for visual extinction and hemianopia using clinical confrontation testing, as well as a perimetry and attention screening test presented on a laptop (participants viewed a central fixation cross while a single gray dot was presented for 66.7ms at one out of 16 possible locations on the screen, for which they indicated via button press if it was perceived). Patients with right hemispheric lesions were also tested for spatial neglect with the Bells Test (Gauthier et al., 1989), the letter cancellation task (Weintraub & Mesulam, 1985), a line bisection task (Ferber & Karnath, 2001) and a copying task (Johannsen & Karnath, 2004). For the neglect tests we used the cut-off scores of CoC = 0.081 for the Bells Test and the CoC = 0.083 for the letter cancellation task (Rorden & Karnath, 2010) and a deviation of 7.77% from the center in the line bisection task (Sperber & Karnath, 2016). The remaining cut-off scores were taken from the test-specific manuals. Participants who showed spatial neglect, hemianopia, clinical object perception deficits or simultanagnosia were excluded from the study. All remaining participants performed without errors or below the cut-off scores for these tests. For the included participants demographic details and results from the VOSP are summarized in Table 2.

**Table 2.**
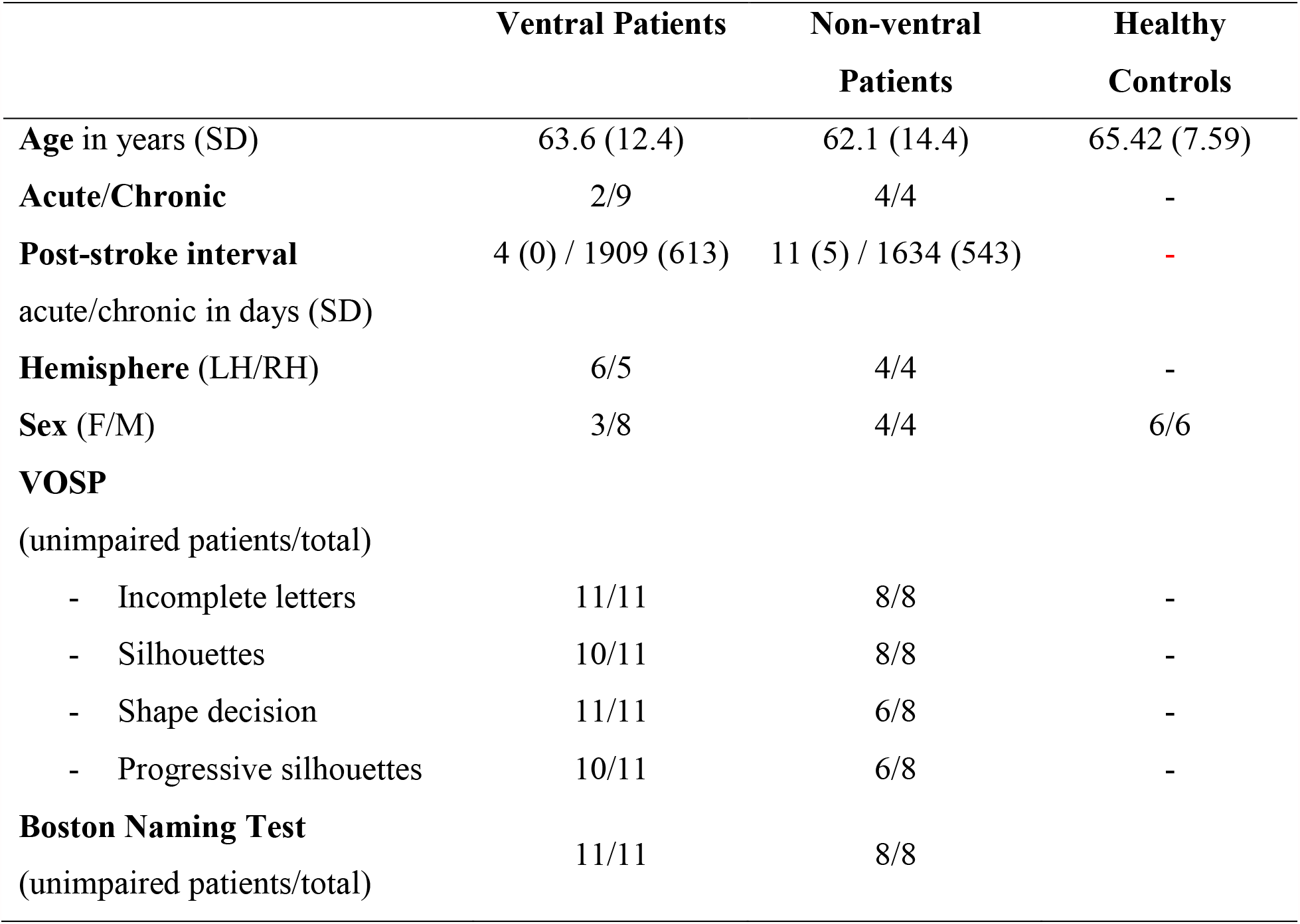
Demographic and clinical details of stroke patients and of healthy controls. The table shows the mean and standard deviations of demographic data and the results of the clinical examination for each subgroup.

|  | Ventral Patients | Non-ventral Patients | Healthy Controls |
| --- | --- | --- | --- |
| **Age** in years (SD) | 63.6 (12.4) | 62.1 (14.4) | 65.42 (7.59) |
| **Acute/Chronic** | 2/9 | 4/4 | - |
| **Post-stroke interval** | 4 (0) / 1909 (613) | 11 (5) / 1634 (543) | - |
| acute/chronic in days (SD) |  |  |  |
| **Hemisphere** (LH/RH) | 6/5 | 4/4 | - |
| **Sex** (F/M) | 3/8 | 4/4 | 6/6 |
| **VOSP** |  |  |  |
| (unimpaired patients/total) |  |  |  |
| - Incomplete letters | 11/11 | 8/8 | - |
| - Silhouettes | 10/11 | 8/8 | - |
| - Shape decision | 11/11 | 6/8 | - |
| - Progressive silhouettes | 10/11 | 6/8 | - |
| **Boston Naming Test** |  |  |  |
| (unimpaired patients/total) | 11/11 | 8/8 |  |

Participants were presented with unilateral grayscale objects either in canonical or non-canonical orientation at the central y-coordinate of a laptop screen with a screen size of 38 cm × 21.5 cm using Matlab (MATLAB, 2016) and the Psychophysics Toolbox (Brainard, 1997). The patient study was conducted with a different set of canonical and non-canonical object stimuli as used in the fMRI-experiment. While stimuli came from the same stimulus set (see above), canonical and non-canonical orientations were determined in an online survey. In a sample of 19 healthy participants (mean age [SD]: 26.84 [8.05]) we asked participants to order three different versions of 49 pre-selected stimuli from their most typical to their most atypical orientation. For each object, the set of images contained always at least one canonical orientation, which was consistent with the definition applied in our fMRI experiment. For the experiment, we selected the most typically (*canonical*) and the most atypically rated version (*non-canonical*). This resulted in a slightly different stimulus set compared to the fMRI study, for which we as described above created frontal and in-depth rotated versions of both canonical and non-canonical views in a different approach. To avoid memory effects the resulting sample was then split into two subsets of images and participants were presented with either only canonical orientations from the first and non-canonical orientations from the second subset or vice versa. Participants were instructed to fixate a central fixation cross while stimuli were presented unilaterally shifted by a visual angle of 11.2° from the center to the left or right at a size of 10.0° visual angle. In six ventral patients with quadrantanopia, experimental stimuli were shifted up (n = 3) or down (n = 3) by 5.4° degrees visual angle to the intact upper or lower visual field. After stimulus presentation, participants named the perceived object and answers were coded by the experimenter. No feedback was given during the experiment to avoid a bias on the results. Each participant conducted two blocks of this task containing images of 14 canonically and 14 non-canonically oriented objects, which were each repeated twice. Stimuli were presented for 83.3ms.

Data was analyzed by fitting linear mixed effects models using the *lmer* function provided by *lme4* package (Bates et al., 2015; *R* Development Core Team, 2018) in *R*. A random intercept for the variable *participant* was included into the model, as well as a random intercept for the variable *trial*, which accounted for the fact that objects were presented twice and a potentially better performance in the second repetition could have occurred. The variables *group* (ventral patients, non-ventral patients, healthy controls), *hemisphere* (left vs. right hemisphere stroke), *hemifield* (ipsi-vs. contralesional stimulus presentation) as well as *view* (canonical, non-canonical) were included as fixed effects. Statistical significance of parameter estimates was assessed using the *Anova* function provided by the *car* package (Fox & Weisberg, 2011; *R* Development Core Team, 2018).

Additionally, we explored the relation between ventral patients’ performance and lesion location. For this purpose, we created reference ROIs of the inferior occipital, the fusiform and the parahippocampal cortex from the AAL atlas (Rolls et al., 2020). We included the amount of overlap of these regions with participants’ lesions as continuous fixed effects in three separate linear mixed effects models, one for each ROI. Thus, the model included the fixed effects *lesion overlap, hemisphere, hemifield* and *view*, as well as all interactive terms. As above, *participant* and *trial* were set as random effects.

## Results

### A. fMRI study

#### Behavioral Data

During the first fMRI experiment, button presses for the task of categorizing objects as made from metal or not were only recorded in 84% of trials for the first eleven participants due to technical problems. For the remaining 9 participants without technical problems, answers were recorded in 93% of trials. We observed that participants classified objects significantly more correctly (*χ* ^2^ = 5.14, *p* = 0.023) for canonically (84%) than for non-canonically presented views (81%). In the group *without* technical problems participants classified objects in canonical views correctly in 87% of cases and objects in non-canonical views in 82% of cases (*z* = -3.263, *p* = 0.001). We did not observe significant differences in the accuracy of frontal and in-depth rotated objects (*p* > 0.097).

#### Univariate ROI Analysis

We statistically compared mean percent signal change values between conditions in the three different ROIs by calculating linear mixed-effects models. Fixed effects were included for *orientation* (canonical *vs*. non-canonical orientation), *rotation* (in-depth rotation *vs*. frontal views) and *hemisphere* (left *vs*. right). *Participant* was included as a random effect.

##### ‘Objects ROI’

We observed a significant main effect for *orientation* (canonical views: 0.85% non-canonical views: 1.04%; *χ*^2^ = 44.665, *p* = 2.34 × 10^−11^) and for *rotation* (in-depth rotated: 0.98%, frontal views: 0.90%; *χ*^2^ = 7.042, *p* = 0.008), but not for *hemisphere* (left hemisphere: 0.93%, right hemisphere: 0.96%; *χ* ^2^ = 0.850, *p* = 0.356). No interactions were significant (full model output: Table 3; estimated marginal means: Figure 4).

**Table 3.**
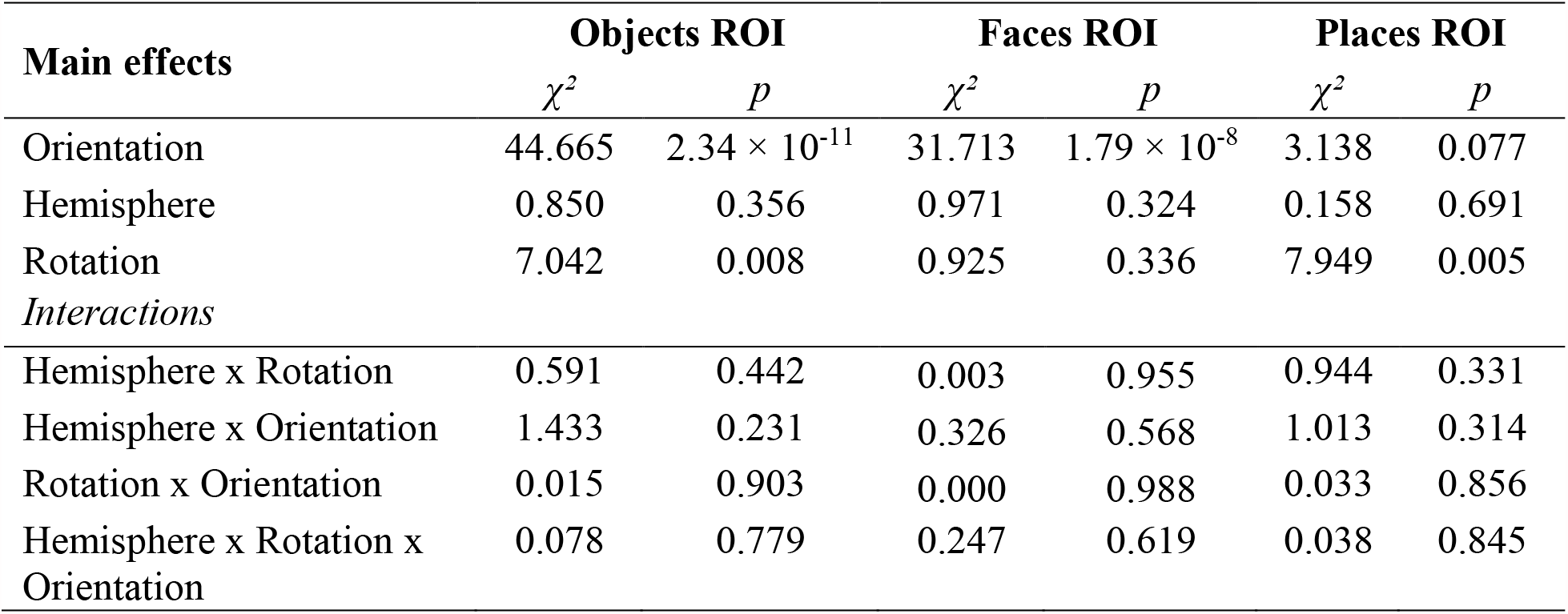
Results of the univariate ROI analyses. We extracted the mean percent signal change values of canonical and non-canonical objects in frontal and in-depth rotated views from individual objects, faces and places ROIs. *Orientation, hemisphere* and *rotation* were set as fixed effects and *participant* was set as random effects in three separate linear mixed-effects models. The table shows the results of the chi-square test of the model, with one degree of freedom for each factor.

**Figure 4.**
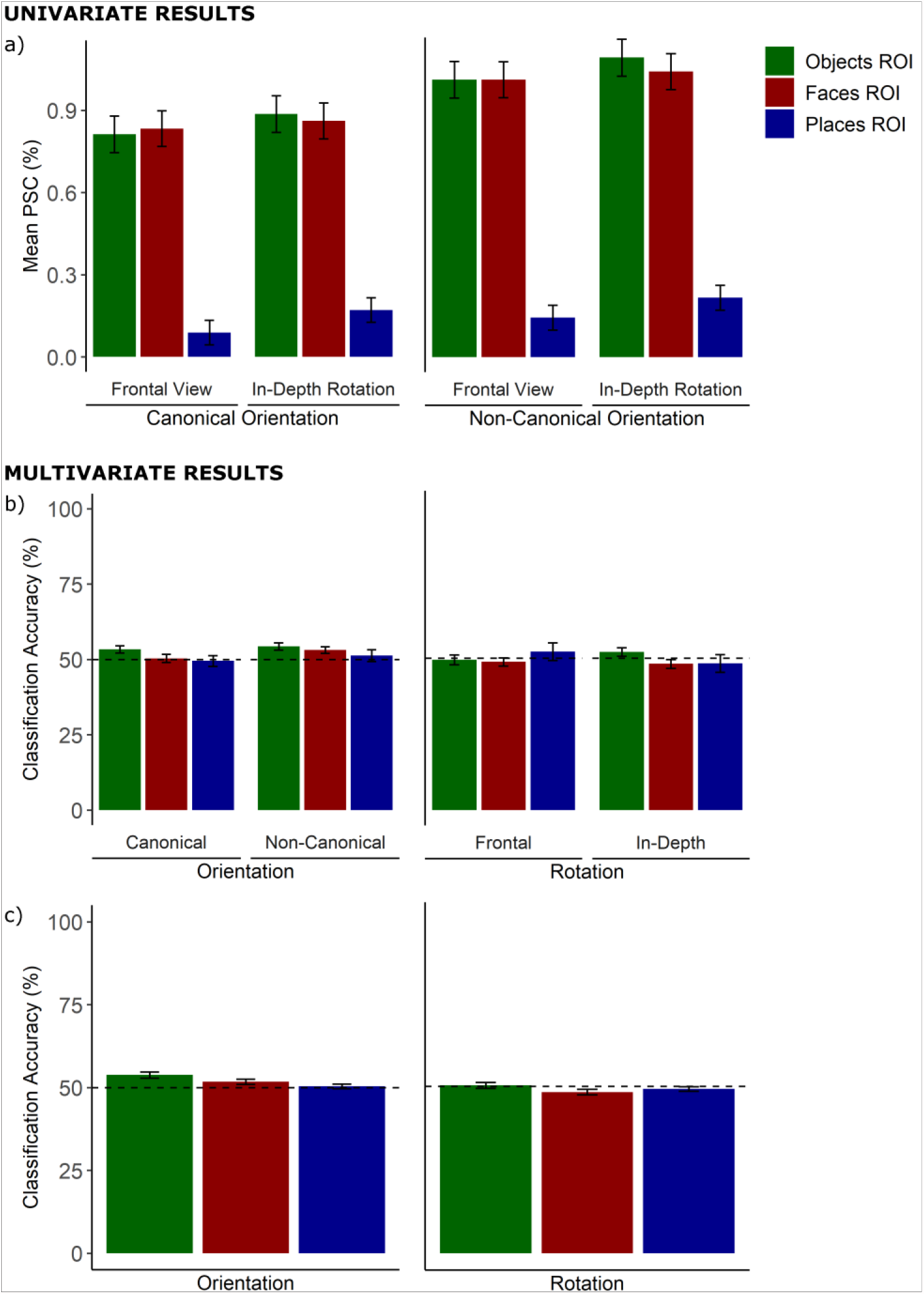
Results of the fMRI experiment. a) Estimated marginal means with error bars (standard error of the mean) from the three ventral ROIs for *canonical* and *non-canonical*/*frontal* and *in-depth* rotated objects. b) Average predictive values (calculated as the proportion of correctly classified trials relative to all tested trials for both object viewing conditions) from the three ventral ROIs for *canonical* and *non-canonical*/*frontal* and *in-depth* rotated objects. c) overall predictive values for *orientation* and *rotation* in the three different ROIs. The dashed line indicates the 50% chance level.

##### ‘Faces ROI’

We observed a significant main effect for *orientation* (canonical views: 0.85% non-canonical views: 1.02%; *χ* ^2^ = 31.713, *p* = 1.787 × 10^−8^) but not for *rotation* (in-depth rotated: 0.95%, frontal views: 0.92%; *χ* ^2^ = 0.925, *p* = 0.336) or *hemisphere* (left hemisphere: 0.95%, right hemisphere: 0.92%; *χ* ^2^ = 0.971, *p* = 0.324). No interactions were significant (full model output: Table 3; estimated marginal means: Figure 4).

##### ‘Places ROI’

We observed a significant main effect for *rotation* (in-depth rotated: 0.19%, frontal views: 0.12%; *χ* ^2^ = 7.949, *p* = 0.005) but not for *orientation* (canonical views: 0.13% non-canonical views: 0.18%; *χ* ^2^ = 3.138, *p* = 0.077) or *hemisphere* (left hemisphere: 0.15%, right hemisphere: 0.16 %; *χ* ^2^ = 0.158, *p* = 0.691). No interactions were significant (full model output: Table 3; estimated marginal means: Figure 4).

#### MVPA

Across all three ROIs, we tested classification accuracies between the left and right hemisphere. We did not find any significant differences (*p* > 0.080) and did not include the factor hemisphere in any of the following analyses. All of the following *t*-tests results represent *t*-tests against 50% chance level. Results of predictive values per condition (*canonical*/*non-canonical* views, *frontal*/*in-depth* views) are shown in Figure 3b. The overall classification accuracies (*orientation*/*rotation*) are presented in Figure 3c.

In the ‘Objects ROI’, we observed predictive values above chance for non-canonical objects (mean: 54.3%, *t*_*(39)*_= 3.64, *p* = 0.003) and canonical objects (53.3%, *t*_*(39)*_= 2.76, *p* = 0.027). The overall classification accuracy for *orientation* was significantly above chance level (53.8%, *t*_*(39)*_= 4.17, *p* < 0.001). We did not observe predictive values above chance for *frontal* views (49.4%, *t*_*(39)*_= -0.38, *p* = 0.706) and *in-depth* views (52.0%, *t*_*(39)*_= 1.46, *p* = 0.302). The overall classification accuracy *rotation* was not significantly above chance level (50.3%, *t*_*(39)*_= 0.77, *p* = 0.767).

In the ‘Faces ROI’, predictive values above chance were observed for non-canonical objects (53.1%, *t*_*(39)*_ = 2.82, *p* = 0.030) but not canonical objects (50.3%, *t*_*(39)*_= 0.23, *p* = 0.818). The overall classification accuracy for *orientation* was significantly above chance level (51.7%, *t*_*(39)*_= 2.21, *p* = 0.033). There were no predictive values above chance for *frontal* views (48.8%, *t*_*(39)*_= -0.90, *p* = 0.744) and *in-depth* views (48.1%, *t*_*(39)*_= -1.28, *p* = 0.620). The overall classification accuracy *rotation* was significantly below chance level (48.2%, *t*_*(39)*_ = -2.13, *p* = 0.040).

In the ‘Places ROI’, we did not observe predictive values above chance for non-canonical objects (51.3%, *t*_*(39)*_= 0.67, *p* = 0.768) and canonical objects (49.5%, *t*_*(39)*_= -0.30, *p* = 0. 768). The overall classification accuracy for *orientation* was not above chance level (50.4%, *t*_*(39)*_= 0.58, *p* = 0.564). We did not observe predictive values above chance for *frontal* views (52.1%, *t*_*(39)*_= 0.74, *p* = 0.768) and *in-depth* views (48.3%, *t*_*(39)*_= -0.60, *p* = 0.768). The overall classification accuracy *rotation* was not significantly above chance level (49.2%; *t*_*(39)*_= -1.23, *p* = 0.450).

To assess statistical differences of SVM classification between object viewing conditions and ROIs we used predictive values as a dependent variable in a linear mixed-effects model for *orientation* and one for *rotation*. The model for *orientation* was calculated with fixed effects for *orientation* (canonical *vs*. non-canonical orientation) and *ROI* (objects, faces, places) and *participant* as a random effect. We found a borderline not significant main effect for *ROI* (*χ* ^2^ = 5.85, *p* = 0.053) and no other significant main effects or interactions (*p* > 0.103). For the model for *rotation*, we included fixed effects for *rotation* (frontal *vs*. in-depth) and *ROI* (objects, faces, places) and *participant* as a random effect. We did not observe a significant main effect or interaction (*p* > 0.277).

All *t*-tests of classification accuracies against chance were corrected for multiple comparisons using the *False Discovery Rate (FDR)* as proposed by Benjamini and Hochberg (1995) per ROI.

### B. Patient study

Table 4 shows parameter estimates and the results of the analysis. There was a significant main effect of the factor *group* (*χ* ^2^ = 10.695, *p* = 0.005), a significant main effect of the factor *view* (*χ* ^2^ = 223.53, *p* < 0.001) and a significant interaction of *view* × *hemisphere* (*χ* ^2^ = 7.54, *p* = 0.006). We used the function *emmeans* (Searle et al., 1980) to calculate estimated marginal means as well as to perform pairwise post-hoc tests (supplementary Table S1). Multiple comparisons were accounted for by using Tukey method. Post hoc tests revealed significant differences in performance only between healthy controls and ventral patients (healthy controls: 88.8%, ventral patients: 73.5%; *t* = 3.290, *p* = 0.008), but not between healthy controls and non-ventral patients (non-ventral patients: 80.4%; *t* = 0 1.655, *p* = 0.242) or between non-ventral patients and ventral patients (*t* = 1.324, *p* = 0.395). Post-hoc tests for the interaction *view* × *hemisphere* revealed significantly better performance for canonical than for non-canonical objects both in the left (canonical: 93.3%, non-canonical: 64.5%, *t* = 12.499, *p* < 0.001) and the right hemisphere (canonical: 88.9%, non-canonical: 70.1%, *t* = 7.752, *p* < 0.001). We observed no significant effect of the factor *hemifield*. Results can be inspected in Fig. 5.

**Table 4.** Results of the patient study. We compared performances in the object naming task for canonical and non-canonical views in healthy controls, non-ventral patients and ventral patients. *Group, view, hemisphere* and *hemifield* were set as fixed effects and *participant* was set as random effects in a linear mixed-effects models. The table shows the results of the chi-square test of the model.

| *Main Effects* | $\chi^2$ | *p* |
| --- | --- | --- |
| Group | 10.695 | 0.005 |
| Hemisphere | 0.047 | 0.829 |
| Hemifield | 0.287 | 0.592 |
| View | 223.526 | <0.001 |
| *Interactions* |  |  |
| Group x Hemisphere | 0.723 | 0.697 |
| Group x Hemifield | 3.306 | 0.192 |
| Hemisphere x Hemifield | 0.184 | 0.668 |
| Group x View | 5.367 | 0.068 |
| Hemisphere x View | 7.542 | 0.006 |
| Hemifield x View | 0.421 | 0.516 |
| Group x Hemisphere x Hemifield | 0.675 | 0.713 |
| Group x Hemisphere x View | 3.309 | 0.191 |
| Group x Hemifield x View | 0.660 | 0.719 |
| Hemisphere x Hemifield x View | 0.497 | 0.481 |
| Group x Hemisphere x Hemifield x View | 0.562 | 0.755 |

**Figure 5.**
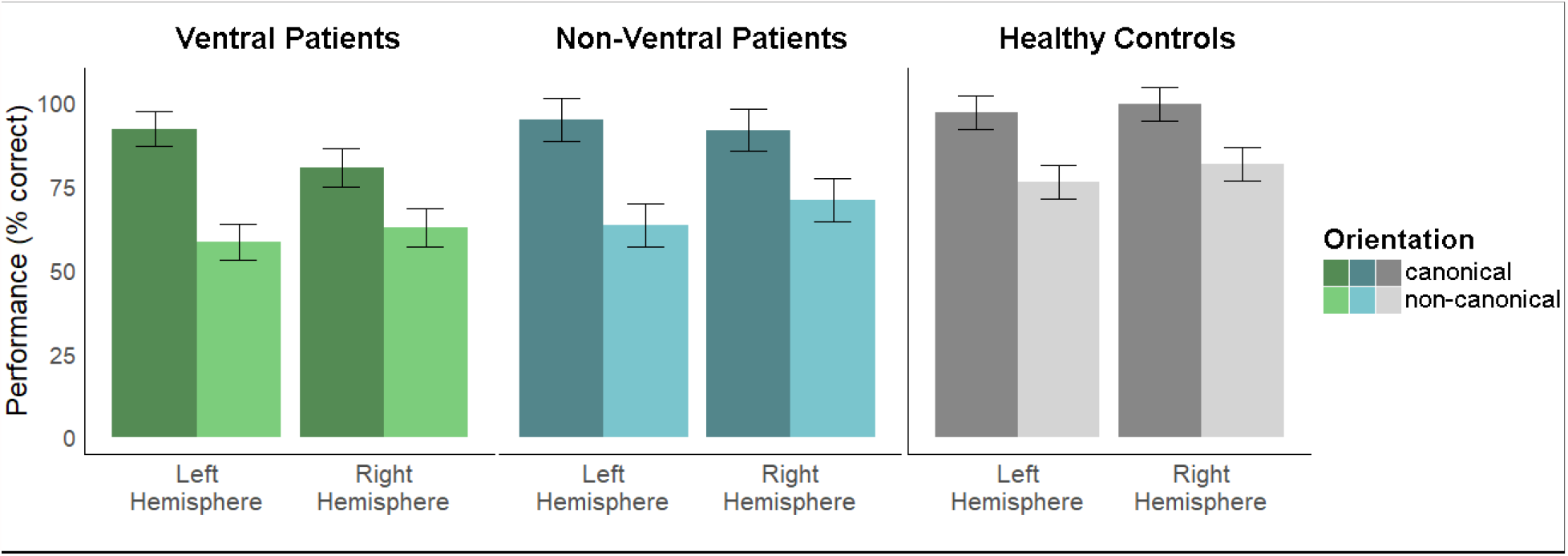
Results of the patient study. Estimated marginal means with error bars (standard error of the mean) from the three subgroups (healthy controls, non-ventral patients, ventral patients) are presented separately for canonical and non-canonical views as well as for lesions of the left and right hemisphere.

The analyses revealed a significant effect of lesion overlap in the fusiform gyrus on task performance (see Table 5 for full model; mere behavioral effects not involving lesion overlap are not shown). We observed significant interactions of the factors *lesion overlap* × *view* (*χ* ^2^ = 7.600, *p* = 0.006), *lesion overlap* × *hemisphere* (*χ* ^2^ = 8.281, *p* = 0.004) and *lesion overlap × hemisphere × view* (*χ* ^2^ = 18.916, *p* = 1.4 × 10^−5^). Post hoc tests revealed significantly different slopes only for lesions of the right fusiform cortex (*t* = 5.048, *p* < 0.001), but not in the left hemisphere (*t* = 1.015, *p* = 0.3136). As indicated by a larger negative estimate (*ß*) for the slope of non-canonical (*ß* = -3.619) than for canonical views (*ß* = -0.797), a larger overlap with the right fusiform cortex might be associated with correspondingly lower performances in naming objects presented in non-canonical views. All other lesion overlap-related effects for the lateral occipito-temporal and parahippocampal cortex were not significant (*p* > 0.106).

**Table 5.** Results of the lesion overlap model in the patient study. We compared the impact of lesion size of the inferior occipital cortex, fusiform gyrus and parahippocampal gyrus on the ventral patients’ performances in the object naming task for canonical and non-canonical views. *Lesion overlap, view, hemisphere* and *hemifield* were set as fixed effects and *participant* was set as random effects in a linear mixed-effects models. As the effects – except of the factor lesion size – were redundant to the analysis conducted before, we report only effects related to the factor *lesion overlap* in order to keep the table clearer. The table shows the results of the chi-square test of the model.

| **Main Effects** | **Inferior**<br>**occipital**<br>**cortex** |  | **Fusiform**<br>**gyrus** |  | **Parahippocampal**<br>**gyrus** |  |
| --- | --- | --- | --- | --- | --- | --- |
| | $\chi^2$ | *p* | $\chi^2$ | *p* | $\chi^2$ | *p* |
| Lesion Overlap | 0.045 | 0.832 | 2.289 | 0.130 | 0.613 | 0.434 |
| **Interactions** |  |  |  |  |  |  |
| Lesion Overlap x Hemisphere | 0.518 | 0.472 | 8.281 | 0.004 | 0.600 | 0.438 |
| Lesion Overlap x Hemifield | 0.001 | 0.970 | 0.754 | 0.385 | 1.843 | 0.175 |
| Lesion Overlap x View | 1.373 | 0.241 | 7.600 | 0.006 | 1.352 | 0.245 |
| Lesion Overlap x Hemisphere x View | 0.043 | 0.835 | 18.916 | <0.001 | 2.608 | 0.106 |
| Lesion Overlap x Hemisphere x Hemifield | 0.031 | 0.859 | 0.108 | 0.743 | 0.103 | 0.748 |
| Lesion Overlap x Hemifield x View | 0.396 | 0.529 | 0.596 | 0.440 | 0.286 | 0.593 |
| Lesion Overlap x Hemisphere x Hemifield x View | 0.455 | 0.500 | 0.054 | 0.817 | 2.507 | 0.113 |

## Discussion

With the present two studies, we aimed at investigating the role of brain areas along the ventral stream in object constancy for object stimuli presented at various viewpoints. In our fMRI experiment with healthy participants, we presented objects in canonical and non-canonical orientations. In anatomically defined ROIs of the lateral occipito-temporal cortex, fusiform and parahippocampal gyri, we selected voxels involved in the processing of objects, faces and places. Within these individual ROIs that were created without spatially normalizing the MRI images, we compared BOLD signals during the presentation of objects in non-canonical and canonical orientation as well as in frontal and in-depth rotated views. We found stronger BOLD activity for objects in non-canonical orientations in the objects and faces ROIs. In-depth rotation was associated with higher activation compared to frontal views in the objects and places ROIs. Applying a machine learning classifier, we demonstrated above-chance classification accuracies for non-canonical and canonical views in the objects ROI and for non-canonical views in the faces ROI. Our second study with stroke patients showed a hemispheric difference where larger lesion overlap with the right hemispheric fusiform gyrus was associated with lower performance in the object recognition task for non-canonical views. Taken together, these results suggest that lateral-occipital and fusiform areas are involved in the processing of objects in non-canonical views and that they might contribute to visual object constancy.

The ventral stream has previously been shown to be organized hierarchically with early stages being involved in the analysis of local elements and later stages in the analysis of complex aspects (e.g., Grill-Spector et al., 1998, 1999; Lerner, 2001; Kravitz et al., 2013; Cowell et al., 2017). According to feature-based theories of object recognition (e.g., Biederman, 1987; Biederman & Gerhardstein, 1993; Chen, 2005), processing of objects in unusual viewing conditions might particularly rely on the detection and analysis of local elements and their integration into holistic percepts (Kosslyn et al., 1994; Lerner, 2001; Epstein et al., 2006). In line with this interpretation, our results suggest that objects and faces ROIs were involved in the processing of local elements in order to identify objects in non-canonical views (Epstein et al. 2006). Higher activations in objects and faces ROIs during the perception of non-canonical compared to canonical views as observed in our fMRI experiment were also found in various previous studies (Kosslyn et al., 1994; Sugio et al., 1999; Haxby et al., 2001; Vuilleumier et al., 2002; Terhune et al., 2005; Nestmann et al., 2021). Epstein et al. (2006) found stronger activations in object sensitive areas in close vicinity of the LOC and middle fusiform cortex in a face and scene inversion experiment. The authors argued that the demanding stimulus presentations evoked activations in brain areas, which are involved in object-processing enabling recognition under demanding viewing conditions. In line with a hierarchical organization of the ventral stream, we did not observe a specific involvement in non-canonical processing in the places ROI, our most anteriorly located ROI. However, our data suggest that this area might be viewpoint-sensitive showing stronger activations for objects, which were rotated in-depth, compared to objects, which were presented in frontal views. Previous studies had shown that the PPA was involved in the processing of objects that are relevant for spatial navigation (Aguirre et al., 1998; Janzen & van Turennout, 2004; Bilalić et al., 2019) with functional connections to the LOC (Baldassano et al., 2013). Our results with higher activations for in-depth rotated views in both objects and places ROIs, suggest that not only spatial processing of scenes but also of object stimuli in the PPA could facilitate complex object perception in earlier ventral stream areas.

Our behavioral study revealed that ventral patients performed significantly worse than non-ventral patients and healthy controls in the object naming task with particularly lower performance for contralesional object presentations. This is in line with recent patient studies showing contralateral object perception deficits after ventral stream lesions (Rennig et al., 2018a,b; Nestmann et al., in press). Moreover, patient and animal studies on viewpoint invariant object perception (Turnbull et al., 1997b) suggested that occipito-temporal brain areas facilitate viewpoint invariant object perception. Beyond, a single-case study reported by Turnbull and McCarthy (1996) demonstrated deficits in the recognition of objects from unusual views after occipital brain injury.

We also observed that patients’ object-naming performance did specifically depend on lesions to the fusiform gyrus. For right hemispheric lesions, our data suggest that larger lesion overlap with the fusiform gyrus was associated with worse performance in the naming of objects in non-canonical views. This result is particularly interesting with respect to a study of de Haan & Campbell (1991) who reported a patient with developmental prosopagnosia, a selective deficit in the recognition of faces, who was also impaired in identifying objects from unusual viewpoints. Prosopagnosia is often reported as a consequence of lesions to the FFA (Cohen et al., 2019) that is commonly associated with face perception (e.g., Sergent et al., 1992; Puce et al., 1995; Kanwisher et al., 1997). In combination with the described research, our results suggest a crucial role of the FFA in processing of objects in demanding viewing conditions. The latter is also in line with results by Vuilleumier et al. (2002) who demonstrated viewpoint-invariant object representations in the left hemispheric fusiform gyrus and sensitivity to object orientation in the right hemispheric fusiform gyrus. In a behavioral experiment with healthy participants, Burgund and Marsolek (2000) assessed viewpoint dependent priming during the presentation of different views of objects in the left or the right hemifield. They observed viewpoint-dependent priming effects only if stimuli were presented in the left visual hemifield, emphasizing viewpoint-specific object perception in the right hemisphere.

Our findings in stroke patients are especially interesting considering previous patient studies that investigated our ability to determine the orientation of objects. Best (1917) reported that a patient with bilateral lesions in the inferior parietal cortex showed unimpaired object recognition but was unable to determine object orientation. Since then, patients with so-called ‘object orientation agnosia’ following such lesion locations have been reported repeatedly (e.g., Harris et al., 2001; Karnath et al., 2000; Turnbull et al., 1997a). Thus, it has been suggested that the orientation of an object is processed to a considerable extent in dorsal stream regions. In line with this, Holmes and Gross (1984) reported that lesions to the ventral path did not affect patients’ ability to discriminate object orientations. However, studies of ‘object orientation agnosia’ have shown a difference between decisions about the upright and non-upright orientation of an object, with the ability to identify the vertical orientation of objects being a more robust function to damage in the dorsal stream area (*cf*. Karnath et al., 2000). Thus, a strict dissociation between the neural representations of object identity and object orientation may fall short. At least some aspects of object orientation appear to be processed outside the dorsal visual stream.

In conclusion, the uni-and multivariate analyses of our fMRI data as well as the results from a parallel patient study argue for a significant contribution of lateral occipito-temporal areas and the (right) fusiform gyrus to object recognition from unusual views. These ventral stream areas might therefore contribute to mechanisms of visual object constancy, possibly functionally different from the processing of object orientation in the dorsal stream.

## Acknowledgements

This work was supported by the Deutsche Forschungsgemeinschaft (RE 3693/1-1 to J.R.). We thank Lisa Röhrig and Daniel Wiesen for assistance during MRI scanning sessions.

## Notes

### Competing Interest Statement

The authors have declared no competing interest.

